# Extracellular Vesicle-delivered Bone Morphogenetic Proteins: A novel paracrine mechanism during embryonic development

**DOI:** 10.1101/321356

**Authors:** Thomas Draebing, Jana Heigwer, Lonny Juergensen, Hugo Albert Katus, David Hassel

## Abstract

Morphogens including Wnt, Hedgehog and BMP proteins are essential during embryonic development and early induction of organ progenitors. Besides free diffusion to form signalling gradients, extracellular vesicle- (EV-) mediated morphogen transport was identified as a central mechanism for Wnt- and Hh-signalling. Here, we investigated EVs isolated from whole zebrafish embryos as a potential morphogen transport mechanism. Inhibition of EV-secretion during development leads to severe dorsalization phenotypes, reminiscent of disrupted BMP-signalling. Subsequently, we found that EVs isolated from zebrafish embryos at bud stage contain biologically active BMP2/4 protein. Embryos with inhibited EV secretion display reduced Smad1/5/9-phosphorylation and downstream gene expression activity. We further show that BMP-containing EVs are secreted by endodermal cells *in vitro*, and inhibition of endodermal-EV release *in vivo* causes signs of BMP signalling loss. Our data provides evidence that establishes the transport of BMP2/4 by EVs as an essential but so far undiscovered mechanism in developmental morphogenesis.

## Introduction

During embryogenesis, a diverse set of cells originates from a single cell of origin. A major task during development is to ensure the correct position and fate of cells. The initially seemingly identical cells start to express a specific set of genes, thereby differentiating into new cell types. This process follows a specific pattern and requires intensive regulation and communication between cells and tissues^1^. Molecules mediating tissue patterning are called morphogens. A hallmark of morphogens is to form a gradient^2^ and to thereby activate concentration dependent signalling responses^3, 4^.

The formation of morphogen gradients is facilitated by multiple components and mechanisms, which are still not fully understood. Extracellular morphogens are typically thought to be secreted into the extracellular space as freely diffusing, unpackaged proteins^5,6^. First models described the gradient formation by diffusion and uptake of morphogens by cells either at the end (‘source-sink’ model^7^) or along the gradient (‘synthesis-diffusion-degradation’ model^7, 8^). More recent findings added more complexity to the existing models. Several soluble and membrane-bound proteins were found to bind morphogens. Depending on the binding factor and context the binding can lead to opposing effects. Thus, binding of bone morphogenetic proteins (BMPs) to Chordin inhibits receptor binding^9, 10^, but can also facilitate increased diffusion rates^11^. BMPs also bind to plasma membrane-bound heparan sulphate proteoglycans (HSPGs), which removes the ability to diffuse freely^12, 13^. Knockout of HSPGs however also disrupts BMP-signalling^12, 13^.

Recently, the morphogens Wnt, TGF-β and Hedgehog have been reported to reside in extracellular vesicles (EVs)^14–17^. The term EV describes a superfamily of vesicles that are characterized by their secretion from cells^18^. EVs include microvesicles (MV) and exosomes among other subtypes. Currently the different types of EVs are mainly characterized by their size, content and cellular compartment, in which they are synthesized. MVs are between 100 – 1.000 nm in size and are formed and shed at the plasma membrane of the secreting cell^19, 20^. Exosomes are the smallest kind of vesicles with a diameter of 50–150 nm in size^19,20^. They form inside multivesicular bodies (MVBs), a part of the endosomal pathway, which can fuse with the plasma membrane, thereby releasing the exosomes^21^. The increasing attention EVs received during recent years is mostly due to their function in transporting specific sets of multiple biomolecules to target cells over long distances^22–25^. The transport of multiple molecules by the same vehicle allows the reliable delivery of complex and fine-tuned signals, while packaging of signalling molecules inside EVs might protect them from degradation and antagonists. Investigating the role of EVs in morphogenetic signalling may thus reveal an additional way of fine tuning morphogenesis during development. Here we report that BMP2 and/or BMP4 (BMP2/4) are secreted in exosome-like EVs *in vitro* and *in vivo* during embryogenesis. This EV-associated BMP was functionally active and activated BMP-dependent transcription in cells. Inhibiting EV-release strongly disturbed BMP-signalling in zebrafish, leading to phenotypes similar to the phenotypes caused by dorsomorphin, a BMP inhibitor. We postulate that EV-association of BMP is an additional tool to shape the BMP gradient.

## Results

### Zebrafish secrete EVs during development

To investigate the role of EVs during embryonic development, we chose the zebrafish as a well-established model in developmental research^26^. EV isolation protocols established so far are mostly optimized to purify EVs from bioliquids, like blood plasma or cell culture media, while only limited options exist for purifying EVs from solid tissue^27,28^. We adapted existing EV isolation procedures to establish a protocol that allowed us an enzyme-free isolation of EVs from early-stage zebrafish embryos by using EDTA and mild shear forces to dissociate embryonic cells, thereby freeing EVs (Figure 1A). EVs were then isolated by sequential ultracentrifugation^14,28^. Nanoparticle tracking analysis (NTA) demonstrated an enrichment of particles with a mode size of 140.3 nm in the isolates, which is within the size range of exosomes (Figure 1B). Electron microscopy showed vesicles of smaller than 200 nm in size, presenting the typical cup-like shape as previously described for EVs^28^ (Figure 1C). Western blot analysis additionally showed the presence of the EV-proteins TSG101, Lamp1 and ALIX in both cell lysate and EV isolates, while the endoplasmic reticulum (ER)-marker GP96 was only present in cell lysate, but not in EVs (Figure 1D). This data indicates a pure EV-isolate with strong enrichment of exosome-like particles secreted by zebrafish embryos.

**Figure 1.**
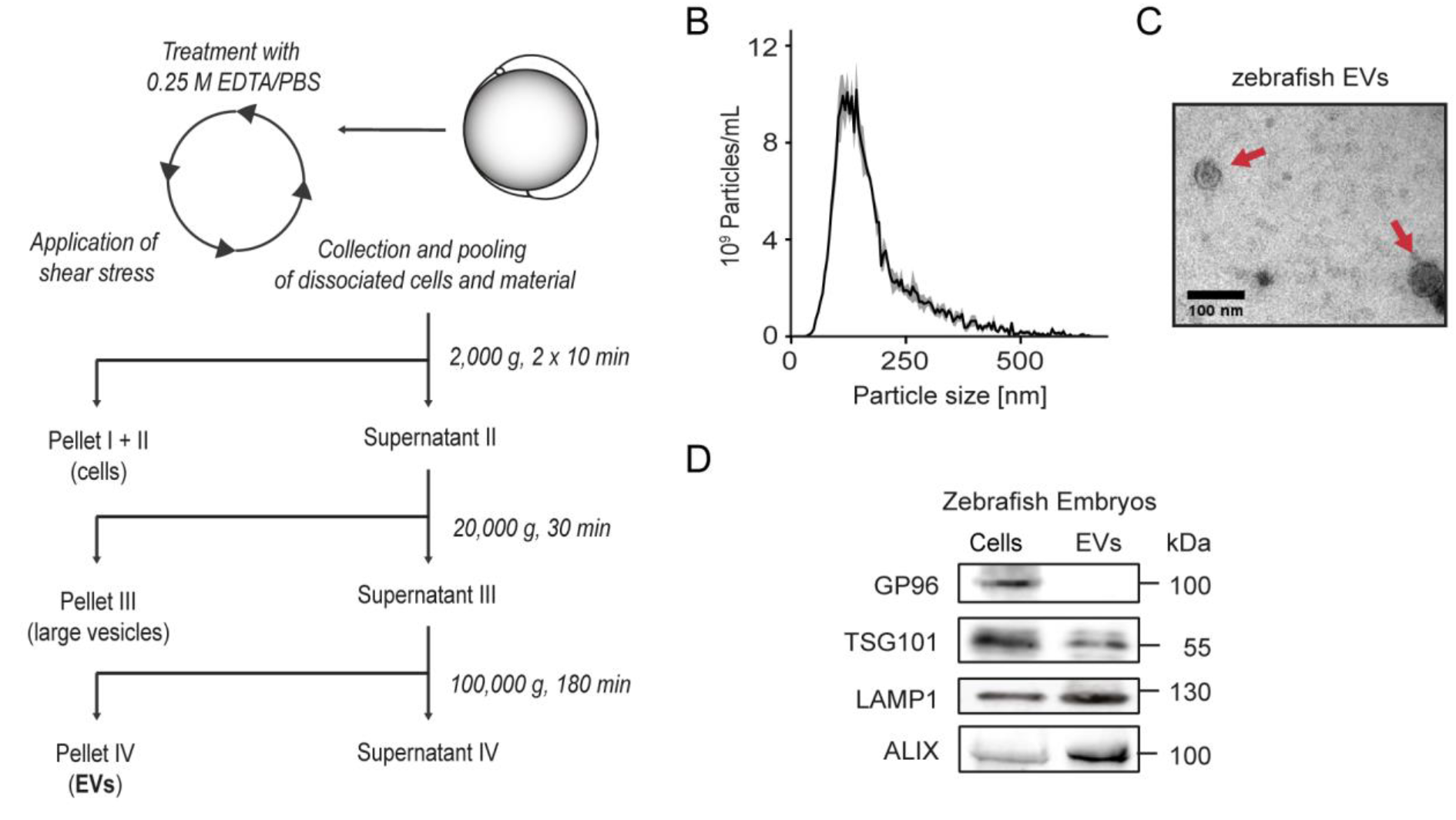
Isolation of EVs from zebrafish embryos. (A) Schematic representation of the protocol used to isolate EVs from zebrafish embryos. Experiments to verify the purity of the EV-isolates were performed with EVs isolated from zebrafish embryos at bud stage (10 hpf): (B) The particle size distribution in the EV-isolates was determined by NTA measurements. The mean of five measurements of the same sample is depicted (black line) with the standard deviation (grey area). (C) The morphology and size of EVs was observed with transmission electron microscopy. Scale bar: 100 nm. (D) Cell lysate (Pellet I/II) and EVs (Pellet IV) from zebrafish embryos were characterized using immunoblots with antibodies detecting GP96, TSG101, LAMP1 and ALIX.

### Inhibition of EV-secretion in zebrafish embryos causes a phenotype resembling dysfunctional BMP-signalling

To evaluate the relevance of EVs during embryonic development, we inhibited EV-secretion by morpholino-based knock down of Rab11a and Rab35. Rab11a and Rab35 are known to be essential for fusion of the multi vesicular body (MVB) with the plasma membrane and therefore are well-established molecular targets to block EV-release into the interstitium^29–31^. To validate the inhibition of EV secretion, EVs were isolated from 130 control- or morphant embryos and compared on the basis of the amount of ALIX present in the samples as determined by western blotting (Figure 2A). EV-isolates of Rab11a-morphants (125 µM MO-Rab11a) showed a non-significant reduction in ALIX content (-14.3 ± 20.7 %) (mean ± standard deviation). Rab35-morphants (250 µM MO-Rab35) however presented a significant reduction of 49.2 ± 29.2 % and were used in further experiments (Figure 2B). The knockdown (KD) of Rab35 resulted in a dorsalization phenotype with varying penetrance and severity (Figure 2C). Classifying phenotype severity using the dorsalization categories published by Mullins et al.^32^ indicated that 34.0 % of the morphants presented severe dorsalization (C4–5), 8.2 % with weak dorsalization (C1–3) and 27.8 % were dead, while only 29.9 % presented no dorsalization phenotype (Figure 2C). Dorsalization is a typical sign of dysfunctional morphogen signalling during dorso-ventral-patterning, especially of reduced BMP2, BMP4 or BMP7-signalling activity^33–36^. Inhibition of BMP signalling by dorsomorphin during embryonic development resulted in an identical phenotype (Figure 2C).

**Figure 2.**
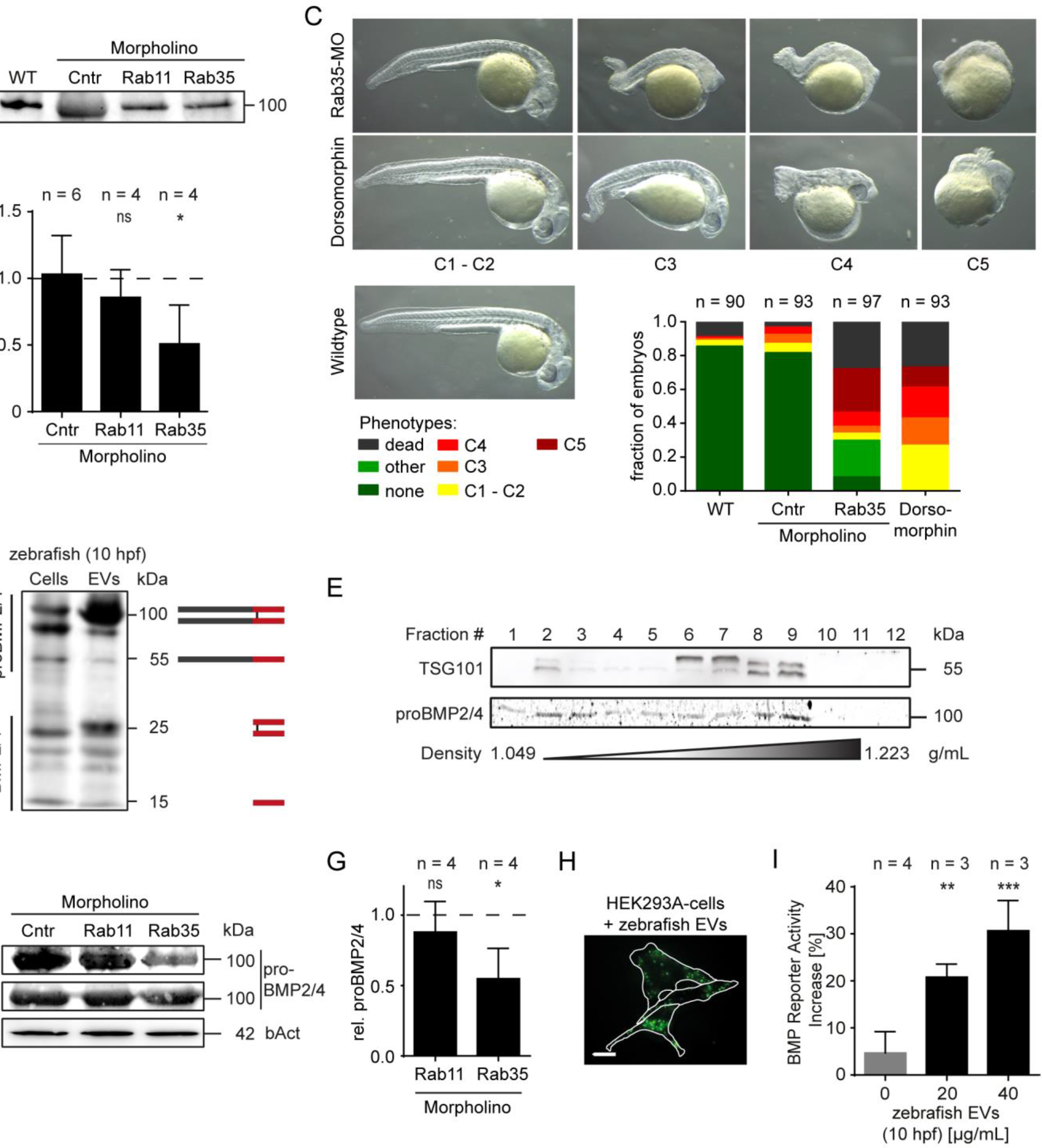
BMP2 and/or BMP4 are transported by EVs. (A) Zebrafish embryos were injected with either 250 µM Cntr-MO, 125 µM Rab11a-MO or 250 µM Rab35-MO at the one-cell stage. EVs were isolated from these zebrafish embryos at bud stage (10 hpf). Western blot analysis with antibodies against ALIX were used to estimate the EV-content of the isolates. The quantification of the western blot signals is plotted in (B). The dashed line represents the measured signal intensity of the wildtype sample, which was used as a normalizer. The samples were compared to the Cntr-MO sample using ANOVA with Tukey’s post hoc test. (C) Images of Rab35-morphants at 24 hpf displaying the spectrum of phenotype severities, resembling dorsalization. For comparison, zebrafish treated with the BMP-signalling inhibitor 10 µM dorsomorphin are shown. The embryos were classified by phenotype severity following the suggestions made by Mullins et al.^32^. The quantification of the fraction of embryos in the respective classes is plotted below the images. (D) EVs were isolated from zebrafish embryos at bud stage (10 hpf) and used in western blot experiments under non-reducing conditions to detect BMP2/4. Multiple preprocessing intermediates of BMP2/4, which are depicted schematically next to the respective protein bands, were detected (dark grey: prodomain; red: ligand domain). (E) EV isolates from zebrafish embryos were further separated on an OptiPrep™-gradient. The gradient was split into 12 fractions, which were investigated by immunoblotting with antibodies targeting TSG101 and BMP2/4. (F) Cell lysate and EVs from Cntr-, Rab11- and Rab35-morphants were investigated for differences in BMP2/4-content by western blot analysis. The quantification of BMP2/4-band intensities in EV isolates is shown in (G). The samples were compared to the normalized mean of the Cntr-morphants (dashed line) using one-sample t-tests with Bonferroni correction for multiple testing. (H) HEK293A-cells were treated with PKH26-labeled EVs isolated from zebrafish embryos and imaged after 5 h. White lines depict cell borders. The scale bar represents 20 µm. (I) HEK293A-cells transiently expressing pGL3-BRE:Luciferase and pIS2-Renilla were treated with unlabelled zebrafish EVs and used in dual luciferase assay 16 h after treatment. Values were normalized to an independent untreated control sample. Samples were compared using ANOVA with Tukey’s post hoc test.

### EVs in zebrafish embryos at bud stage contain BMPs

To assess whether BMPs are transported by embryonic zebrafish EVs, we performed western blotting of zebrafish EVs secreted at bud stage (10 hpf). The antibody was chosen to recognize the mature domain of BMP2, allowing the detection of both processed and unprocessed BMP2. Since the mature domain of BMP2 and BMP4 are nearly identical, the antibody detected both BMPs. Thus, we will refer to the detected BMPs as BMP2 and/or BMP4 (BMP2/4). Surprisingly, we detected the BMP2/4-precursor as well as the mature BMP2/4 ligand (Figure 2D). To confirm that BMP2/4 is associated to EVs and not purified as a contaminant, the crude EV-suspension was separated on an OptiPrep®-gradient (Figure 2E). BMP2/4 mainly co-purified within the same fractions as TSG101 at a density of 1.13 – 1.18 g/mL, which was previously described for EVs^19, 28^. In Rab35-morphants the amount of BMP2/4 in EV-isolates was significantly reduced as compared to EV-isolates obtained from control zebrafish, while BMP2/4-levels remained constant in the cell lysate of the same embryos (Figure 2F,G). Rab11-KD on the other hand did not reduce the amount of BMP2/4 in EV-isolates.

To demonstrate the biological activity of EV-associated BMP, we performed a dual luciferase reporter assay in HEK293A cells using a construct expressing Firefly Luciferase under the control of BMP-responsive elements (BRE)^37^. Since zebrafish EVs might not actually be endocytosed by HEK293A-cells, we verified the uptake by labelling the EVs using PKH26 and imaged EV-treated cells after 5 h (Figure 2H). The cells showed a large number of fluorescent punctae, resembling endosomes, inside their cytosol, thereby indicating EV uptake. HEK293A cells were transfected with pGL3-BRE:Luciferase and pIS2-Renilla and subjected to a 16 h-treatment with either PBS or EVs isolated from zebrafish at bud stage. The EV-treatment resulted in a significant, dose-dependent increase in luciferase activity indicating that EV-delivered BMP indeed is able to activate BMP-dependent transcription (Figure 2I).

### EV-transport of BMP2/4 is required for BMP-signalling during development

To address the hypothesis that BMP signalling is reduced by inhibition of EV-secretion, we used Nkx2.5, a transcriptional target of BMP2/4, as a readout to measure BMP-activated transcriptional activity in Rab35-morphants. Whole mount *in situ* hybridisation (WISH)-stainings of *nkx2.5* at the 7-somite stage (7SS) resulted in a typical staining with two parallel stripes along the midline, representing cardiac progenitor cells, in wildtype zebrafish embryos and control morphants^38^ (Figure 3A). In Rab35-morphants we found the *nkx2.5*-positive areas to be smaller and situated more laterally (Figure 3A). Quantification of the *nkx2.5*-staining showed a significant reduction of the *nkx2.5*-expression to a median of 36.9 % (Quantiles (25 % – 75 %): 12.4–87.8 %) in Rab35-morphants as compared to wildtype zebrafish embryos. These results suggested that inhibition of EV-secretion results in reduced BMP-mediated transcriptional activity.

A well-observable and strong BMP-signalling event during early zebrafish development happens in the tail bud during tail development^39^. We used whole mount immunofluorescence stainings to detect phosphorylated Smad1/5/9 (pSmad1/5/9) in the tail bud of 7SS zebrafish embryos as an additional measure of BMP-signalling activity (Figure 3C). Again, while levels of pSmad1/5/9 were indistinguishable in wildtype and MO-Cntr-injected zebrafish, Rab35-KD significantly reduced the overall Smad1/5/9-phosphorylation in the tail bud to 45.4 % (Quantiles (25 % – 75 %): 28.4–77.0 %) (Figure 3D).

**Figure 3.**
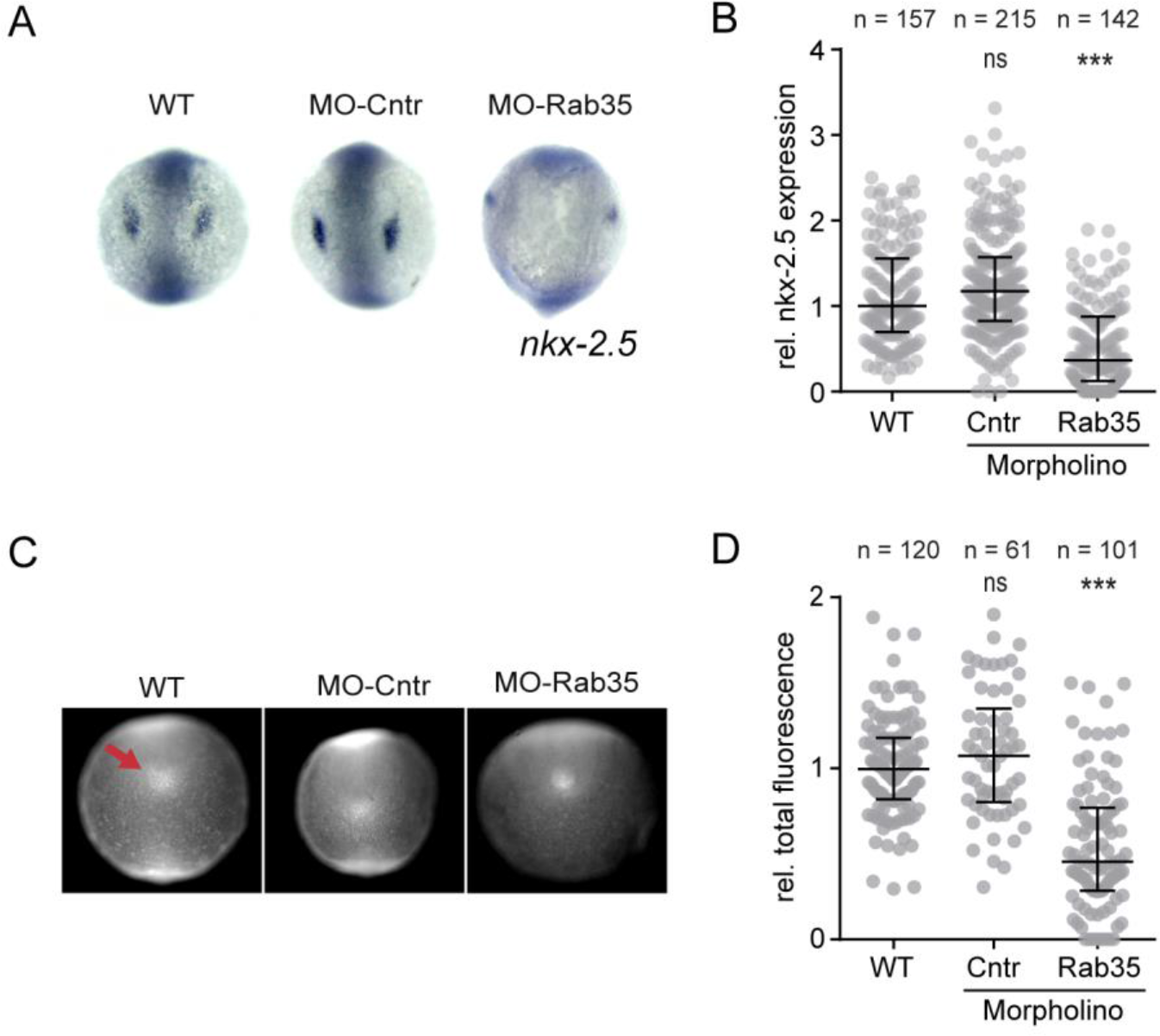
Rab35-KD reduces BMP2/4-dependent signalling activity. Wildtype zebrafish embryos, Cntr- and Rab35-morphants were collected at the 7-somite stage. (A-B) Whole mount *in situ* hybridizations was performed to detect *nkx-2.5*. Image analysis-based quantification was used to measure the *nkx-2.5* expression in the embryos. (C) Typical whole mount immunofluorescence stainings detecting phosphorylated Smad1/5/9 are shown. The tail bud is indicated by the red arrow. (D) The Smad1/5/9-phosphorylation was quantified in the tail bud. The total fluorescence is defined by the product of the stained area, which was normalized to the area of the embryo, and the background corrected fluorescence intensity. All statistical comparisons were performed by using the Kruskal-Wallis-Test with Dunn’s post hoc test. In the plots (C, D) the middle lines represent the median and whiskers represent the 25 % and 75 %-quantiles.

Taken together, these experiments strongly suggest that EVs play a crucial role in mediating BMP signalling activity and that EV-mediated BMP signalling is significantly involved in early cardiac progenitor cell induction.

### Endoderm cells secrete EVs containing BMP2/4

BMP2 responsible for activating the transcription of Nkx2.5 in cardiac mesoderm cells is known to originate from the endoderm^40–42^. Since the EV isolation protocol for zebrafish embryos does not allow purification of EVs originating from a specific tissue or cell type, we used the mouse endodermal cell line End2 as an endodermal *in vitro* model^43^. Using western blotting, we verified the purity of the End2-EVs purified from cell culture medium (Figure 4A). Western blotting also confirmed that BMP2/4 was present in EVs isolated from End2-cells. The purity of End2-EV isolates was further asserted by electron microscopy (Figure 4B) and NTA measurements (Figure 4C), confirming an enrichment of particles smaller than 200 nm with a mode diameter of 140.6 nm. On an OptiPrep®-gradient BMP2/4 largely co-purified in the same fractions as the EV-markers Flot1 and ALIX, suggesting their association (Figure 4D).

**Figure 4.**
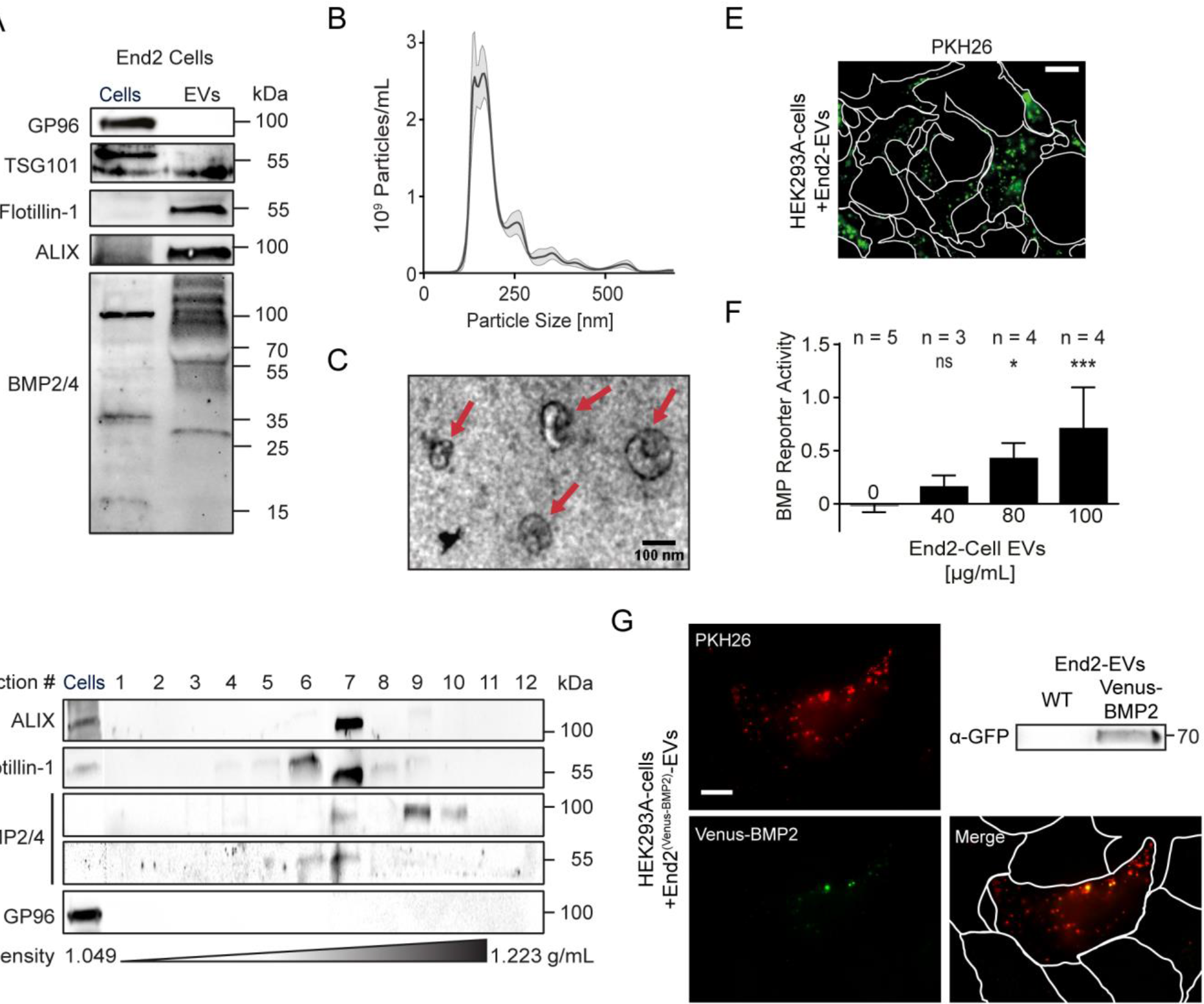
The endoderm is a source for EV-associated BMP2/4. EVs were isolated from End2-cells following the protocol described by Gross et al.^14^. (A) Western blot analysis of End2 cell lysates and End2 EVs using antibodies targeting GP96, TSG101, Flotillin-1, ALIX and BMP2/4. (B) NTA-measurement of a representative End2 EV-isolate. The black line represents the mean of five consecutive measurements. The grey area depicts the standard deviation of these measurements. (C) Representative electron microscopy image of End2-EVs. Red arrows mark exosome-sized EVs. The scale bar represents 100 nm. (D) OptiPrep™-gradient centrifugation was performed on End2-EV isolates. Gradients were separated into 12 fractions and analysed by western blotting. (E) HEK293A-cells were treated with PKH26-labeled End2-EVs for 5 h before life-cell imaging. The scale represents 20 µm. White lines represent cell borders. (F) HEK293A-cells transfected with pGL3-BRE:Luciferase and pIS2-Renilla were treated with End2-EVs for 16 h. BMP-signalling activity was measured using a dual luciferase assay. Values were normalized to an independent untreated control sample. Samples were compared using ANOVA with Tukey’s post hoc test. (G) EVs were isolated from End2-cells transfected with a plasmid encoding Venus-BMP. Western blotting was used to show the presence of Venus-BMP in EV-isolates. HEK293A-cells treated with PKH26-labeled End2^Venus BMP^-EVs for 5 h. The scale bar represents 20 µm. The white lines represent cell borders. Merged image shows co-localization of Venus-BMP and PKH26, indicating association of BMP with EVs.

PKH26-labeling again confirmed the uptake of End2-derived EVs by HEK293A-cells (Figure 4E), providing the basis for dual luciferase assays to test BMP-activity. As expected, the BMP-activity assay verified the ability of End2-EVs to activate BMP-dependent transcription (Figure 4F). Further, fluorescently labelled BMP2 (Venus-BMP2) expressed in End2-cells was detected in EV-isolates. After treatment of HEK293A-cells with PKH26-labelled EVs from Venus-BMP2-expressing End2-cells, overlapping fluorescent signals for PKH26 and Venus were found in the HEK293A-cells, again confirming EV and BMP 2/4 association (Figure 4G).

### Endoderm-specific inhibition of EV-secretion dampens Smad1/5/9-phosphorylation in the zebrafish tail bud ***in vivo***

To assess the significance of the endoderm as a source of EV-associated BMP2/4 *in vivo* we injected zebrafish embryos with a plasmid construct expressing a dominant negative variant of Rab35 (Rab35^N120I^)^29^ tagged with an N-terminal GFP (GFP-Rab35^dn^). An endoderm-specific expression was achieved by using the *sox17*-promoter^44^. Since zebrafish stably expressing GFP-Rab35^dn^ were not viable, transient expression was used instead, leading to a mosaic expression pattern only. Nevertheless, the expression of Rab35^dn^ in the endoderm resulted in a similar phenotype as the ubiquitous Rab35-KD (Figure 5A, S1A), indicating that the endoderm is an important source for EVs involved in BMP-signalling. Surprisingly, overexpressing the wildtype form of Rab35 in the endoderm also increased the mortality and number of dorsalized embryos, although to a lesser extent than Rab35^dn^.

**Figure 5.**
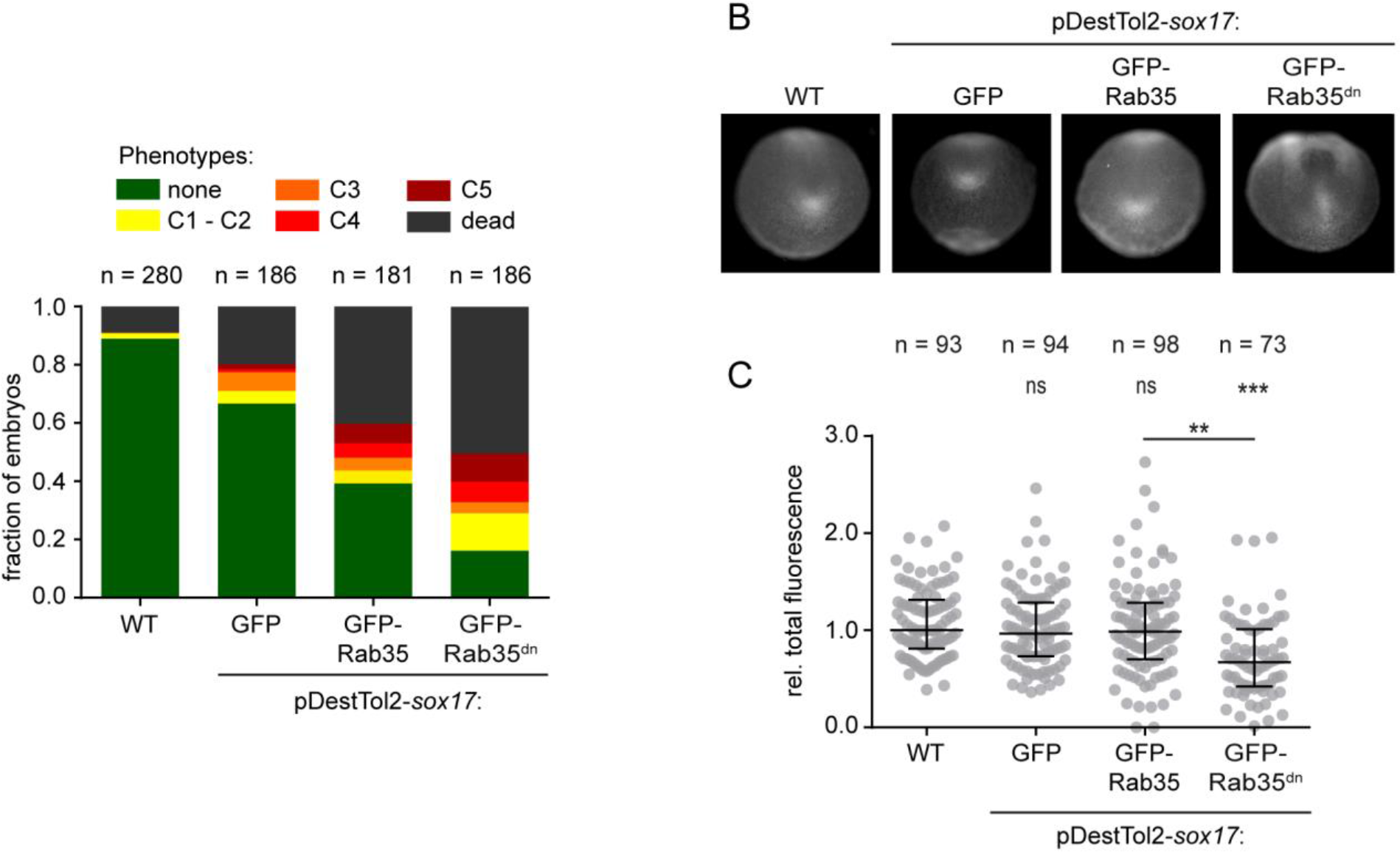
EV-secretion from the endoderm is needed for BMP-signalling during early development. Zebrafish embryos were injected with a pDestTol2 vector encoding either GFP, GFP-Rab35 or the dominant negative GFP-Rab35^N120I(dn)^ under the transcriptional control of the *sox17*-promoter. (A) Phenotypes were assessed at 24 hpf. (B) Whole mount immunofluorescence stainings were used to label pSmad1/5/9. Embryos were imaged in the posterior view. (C) The pSmad1/5/9 labelling intensity in the tail bud was quantified. The distributions were compared using the Kruskal-Wallis-test in combination with Dunn’s post hoc test.

Immune fluorescence stainings against pSmad1/5/9 showed that the expression of GFP-Rab35^dn^ in contrast to expression of GFP or GFP-Rab35 resulted in morphological changes and reduced fluorescence in the tail bud (Figure 5B). Quantification of the pSmad1/5/9 distribution and intensity in the tail bud validated these initial observations (Figure 5C). Interestingly, no overlap of the transgene-expressing cells with cells positive for pSmad1/5/9 was needed to cause the observed phenotype, indicating that a paracrine effect might be the cause for the reduced Smad1/5/9-phosphorylation (Figure S1C). Thus, our results suggest that endoderm-derived EVs are required for BMP-signalling during zebrafish development.

### BMP2/4 is tethered to EV surfaces by binding to HSPGs

BMPs, other than Wnts or Hh, do not possess a lipid anchor, with which they can bind to EV-membranes^45–47^. However, BMPs are known to bind to heparan sulphate proteoglycans (HSPGs) on cell surfaces ^13, 48, 49^. Since HSPGs are present on EV-surfaces as well^50^, we investigated whether BMPs are tethered to EVs by HSPGs. We divided End2-EV isolates into two equal fractions. Fraction I was treated with Heparinase 3, which cleaves off the sugar chains of HSPGs to which BMPs are bound^51^. Fraction II was used as a control. If HSPGs are present and are responsible in linking BMPs to EVs, Heparinase 3 treatment should result in the decrease in BMP signal using western blot. Heparinase 3-treatment indeed led to a decrease in BMP2/4 in the EV-isolate (Figure 6A,B), indicating that BMP2/4 is tethered to the EV surface in part by binding to HSPGs.

**Figure 6.**
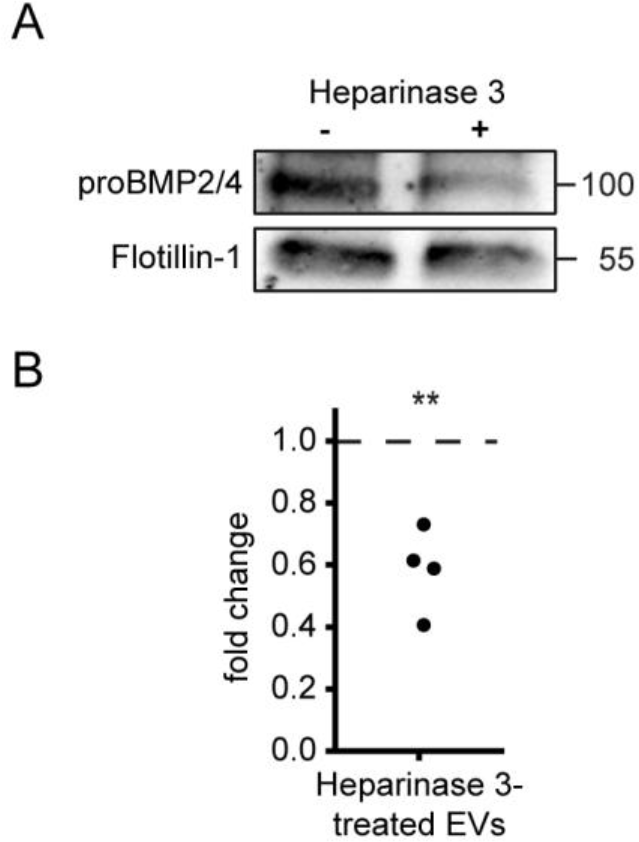
BMP2/4 is tethered to EV-surfaces by binding to HSPGs. End2-EV isolates were split into two equal fractions and either treated with Heparinase 3 or with Heparinase 3 reaction buffer. (A) Representative immunoblot of Heparinase 3-treated and control EV-samples. (B) proBMP2/4-band intensities were normalized with the respective Flotillin-1-band intensities. The Heparinase 3-treated sample was further normalized to the control and a one-sample t-test was used to determine whether the amount of proBMP2/4 in EVs was significantly reduced after Heparinase 3-treatment.

## Discussion

Here, we report that BMP2 and/or BMP4 are transported by EVs during zebrafish development, a mechanism that was essential for BMP-dependent morphogenetic signalling during zebrafish development. We further provide a mechanism of tethering morphogens without lipid-modification to EVs.

Morphogenetic signalling has been investigated for over a century and while it is commonly accepted that diffusion gradients are a hallmark feature of morphogen signalling, the mechanisms behind the formation of morphogen gradients are still not definitely resolved^8, 52^. Differences in morphogen gradient formation do not only occur between different morphogens, but also between morphogenetic signalling events facilitated by the same morphogen. The Dpp-gradient during the dorsoventral-patterning in drosophila, for example, is formed within 30 min by an active shuttling mechanism involving Sog^53,54^. In the imaginal wing disc on the other hand, the Dpp-gradient needs 4 h to form, which is thought to happen by a different unaided mechanism^55^. Several additional models were proposed based on experimental observations. This includes the Source-Sink-model^7,56,57^, the Counter-Gradient-model^56–58^, restricted diffusion^11,12^, transcytosis^59^, cytoneme transport^60^ and more recently also vesicular transport^14–16, 61^. This role of EVs in morphogenetic signalling is by now a well-established mechanism after Wnt-proteins and Hh were found in EVs, a mechanism that allows these lipid-modified proteins to diffuse efficiently without unspecifically binding to membranes^14–16^. The observation that BMPs, that do not possess a lipid anchor, are transported by EVs as well, indicates that masking lipid groups is not the only function of EVs in morphogenetic signalling. EVs might be needed to mobilize BMP that is bound to HSPGs on cell surfaces and thus restricted in their diffusion.

Multiple proteomics studies detected BMP in EVs but did not investigate this finding in detail^62–69^. Furthermore, the secretion of BMPs in matrix vesicles, a specialized type of EV that is involved in bone formation was reported earlier by Nahar et al.^61^, further supporting our findings.

The phenotypical changes observed in zebrafish embryos after Rab35-KD suggest that EV-transport is essential for BMP-signalling during embryonic development. The severity of the phenotypes is surprising, considering that BMP is thought to diffuse unaided. But whether the importance of EVs in BMP-signalling is due to their ability to mobilize BMPs or due to other reasons, remains to be investigated. A limitation of this study is that known factors involved in EV-secretion, that can be manipulated to inhibit the release of EVs, are also involved in essential cellular processes, especially endosomal trafficking, including receptor recycling^29,70,71^. Thus, unspecific effects of changes in Rab35-activity might add to the observed phenotype. Our experiments however show that inhibition of Rab35 in the endoderm affected BMP-signalling activity in cells of the zebrafish tailbud, which are mesoderm cells^72^. Processes that take part in computing the BMP-signal response in the mesodermal target cells, as for example BMP-receptor recycling, cannot be influenced by Rab35^dn^ directly, that is exclusively expressed in the endoderm. This indicates, that a paracrine effect is likely responsible for the reduced Smad1/5/9-phosphorylation.

The finding, that EV-associated BMP2/4 is secreted by the endoderm, is intuitive, since the endoderm was already described to be responsible for secreting freely diffusing BMP2/4 during early development^40–42^. Nevertheless, since BMP2/4 associate to EVs by binding to HSPGs on their surface should theoretically be able to catch free BMP2/4 in the extracellular space. This would mean that EVs from cells, which are not secreting BMP2/4, could be able to influence BMP2/4-signaling by secreting BMP-tethering EVs, thereby modulating BMP gradient formation and thus BMP signalling. Whether this hypothesis is true, remains to be investigated in future studies.

In summary, we describe an additional mode of transport for BMPs during embryonic development. While future studies will have to show how EV-associated BMPs contribute to BMP-signalling as compared to previously known mechanisms, the findings presented here provide additional information, that will help to understand the formation of BMP-gradients.

## Experimental Procedures

### Zebrafish care and maintenance

All animal experiments have been performed in accordance with the guidelines of the state of Baden-Wuerttemberg and all experimental protocols. Wildtype zebrafish were used as basis for all experiments. For imaging, zebrafish embryos were mounted in 2% methyl-cellulose and analysed under the microscope.

### Inhibition of EV secretion

Morpholinos were obtained from Gene Tools LLC. The morpholinos against Rab11 and Rab35 were designed to bind at the translation start site. The sequences were as follows: Rab11^73^: 5’-GTATTCGTCGTCTCGTGTCCCCATC-3’; Rab35^74^: 5’-AGAGGTGATCGTAGTCGCGGGCCAT-3’. MO-Cntr: non-targeting standard control (Gene Tools LLC). Morpholinos were solubilized in water at a stock-concentration of 1 mM. Stock solutions were diluted with 200 mM KCl to working concentrations of 125 μM–250 μM. The injection volume was 2–4 nL/embryo. Effective doses were determined for every morpholino separately.

Inhibition of EV secretion was verified by western blot analysis using an antibody targeting ALIX (see below). EVs of uninjected and MO-Cntr-injected zebrafish embryos were used as controls. Experiments showing large differences in ALIX-content of EVs between uninjected and MO-Cntr-injected zebrafish embryos were excluded.

### Cell Culture

HEK293A-cells were maintained in Dulbecco’s Modified Eagle Medium (DMEM) (Gibco) containing 10 % Foetal Calf Serum (FCS) (Gibco), 1 % L-Glutamine (Gibco) and 1 % Penicillin/Streptavidin (Gibco). End2 cells^43^ were maintained in DMEM/F-12 medium (Gibco) containing 7.5 % FCS and 1 % Penicillin/Streptavidin. All cells were cultured at 37 °C and 5 % CO_2_ in a humid atmosphere. Transient transfections of plasmids were performed using Lipofectamine 2000 (Invitrogen) following the manufacturer’s instructions.

### EV Purification of Cell Culture Medium

Cells were grown to 70 % confluency. The medium was exchanged with fresh medium containing FCS, that was depleted from EVs by centrifugation at 100,000 g for 18 h and successively filtered through a 0.2 µm filter. After 24 h the conditioned medium was collected and EVs were isolated by ultracentrifugation following the protocol described by Gross et al.^14^.

### Nanoparticle Tracking Analysis

The EV pellet were resuspended in PBS. Particle counts and size distribution were measured using a NanoSight LM10 (Malvern Instruments) equipped with a 405 nm laser.

### Electron Microscopy

EVs resuspended in PBS were prepared for electron microscopy following the protocol described by Théry et al.^28^

### OptiPrep Gradient Centrifugation

EV pellets were resuspended in 500 µL Homogenization Media (0.25 M sucrose, 10 mM Tris-HCL, (pH 7.4)). 40 %, 20 %, 10 % and 5 % OptiPrep/Homogenization Media solutions were prepared. The gradient was created by layering 3 mL of the 40 %, 20 % and 10 % OptiPrep solutions and 2.5 mL of the 5 % OptiPrep solution in order. The EV sample was layered on top of the gradient. The gradients were ultracentrifuged at 100,000 g for 18 h at 4 °C. Fractions of 1 mL were collected beginning from the top of the gradient and diluted with PBS. EVs in each fraction were isolated by ultracentrifugation at 100,000 g for 3 h at 4 °C.

### Imaging EV uptake

For labelling with PKH26 (Sigma Aldrich) the EV pellet was resuspended in 1 mL Diluent C. 2 µL of PKH26 was added and the samples were incubated for 30 min at room temperature. Afterwards the sample was diluted by adding 11 mL of PBS. The labelled EVs were collected by ultracentrifugation at 100,000 g for 3 h at 4 °C. The EV pellet was resuspended in PBS. Cells were treated with 50 ug/mL of PKH26-labelled EVs. Life-cell imaging was performed 5 h after treatment.

### ***In vitro*** BMP activity assay

HEK293A cells seeded in 96-well plates were transfected with 10 ng pIS2-Renilla Control Vector and 100 ng pGL3-BRE:Luciferase^37^. The medium was exchanged to serum-free medium 1 h before treatment. Cells were either treated with EVs or PBS. 16 h post treatment cells were analysed using the Dual Luciferase Reporter Assay system (Promega, Cat.-No. E1960) following the manufacturer’s instructions. Renilla Luciferase activity was used to normalize Firefly Luciferase activity.

### Whole mount in situ hybridization

The whole mount in situ hybridization was performed as previously described by Thisse et al.^75^ The probes to detect nkx2.5 were generated using the primer pair 5’-CGGGACATACTGAACCTGGA-3’, 5’-TCTCCCAGACACGGTTTACC-3’. Staining intensity was quantified using Fiji (ImageJ). Images were contrasted consistently in each experiment. Each stained area (both heart fields) was segmented manually. The area of the staining was normalized to the total area of the embryo. The background intensity was measured close to the stained area and used for background subtraction. The *nkx2.5*-expression was defined as the product of the background-corrected mean staining intensity and the normalized staining area.

### Whole mount immunofluorescence

Zebrafish embryos fixed in 4 % PFA were dehydrated with an ascending methanol series. Before staining, the embryos were rehydrated with a descending methanol series and permeabilized by incubation with ice cold acetone for 8 min at –20 °C. Embryos were blocked in PBS with 1 % Triton-X100, 2 % BSA, 10 % sheep serum and 1 % DMSO for 30 min before incubation with the primary antibody for at least 20 h at 4 °C. Incubation with the secondary antibody was performed overnight at 4 °C. The staining intensity in the tail bud of the embryos was analysed in the same way as the *nkx2.5*-whole mount *in situ* stainings.

### Heparinase 3-treatment

EVs resuspended in PBS were divided into two equal parts of which one was treated with 2 mIU/mL Heparinase 3 in reaction buffer (20 mM TrisHCl, 0.1 mg/mL BSA, 4 mM CaCl_2_ (pH 7.5)) and the other with reaction buffer for 3 h at 37 °C. EVs were isolated from samples by ultracentrifugation with 100,000 g for 3 h at 4 °C before further analysis.

### Antibodies

For western blotting experiments the following antibodies were used: GP96 (CST, #2104), TSG101 (Sigma Aldrich, AV38773), ALIX (Sigma Aldrich, sab4200476), Flotillin- 1 (BD Biosciences, 610821), LAMP1 (Sigma Aldrich, sab3500285), beta-Actin (Sigma Aldrich, A5441), pSmad1/5/9 (CST, #13820) and BMP2/4 (RnD Systems, MAB1128).

### Statistics

Statistical tests used for analysis are indicated in the respective figure captions. Significance levels are indicated as follows: ns: not significant; *: p ≤ 0.05; **: p ≤ 0.01; ***: p ≤ 0.001. If not indicated otherwise error bars represent the standard deviation.

## Acknowledgements

We thank Prof. Dr. Michael Boutros for allowing us to perform NTA-measurements at his lab, Ms. Hosser for taking the electron microscopy images, Prof. Dr. Mummery for providing us with the End2-cell line, Prof. Dr. Mikael Simons for sharing Rab35-constructs and Prof. Dr. De Robertis for the Venus-BMP2 construct.

## Author Contributions

D.H. and T.D. designed the experiments and wrote the article. T.D., J.H. and L.J. performed the experiments. T.D. analysed the data. H.A.K. and D.H. provided the material, read and revised the article.

## Supplementary Information

**Figure S1.**
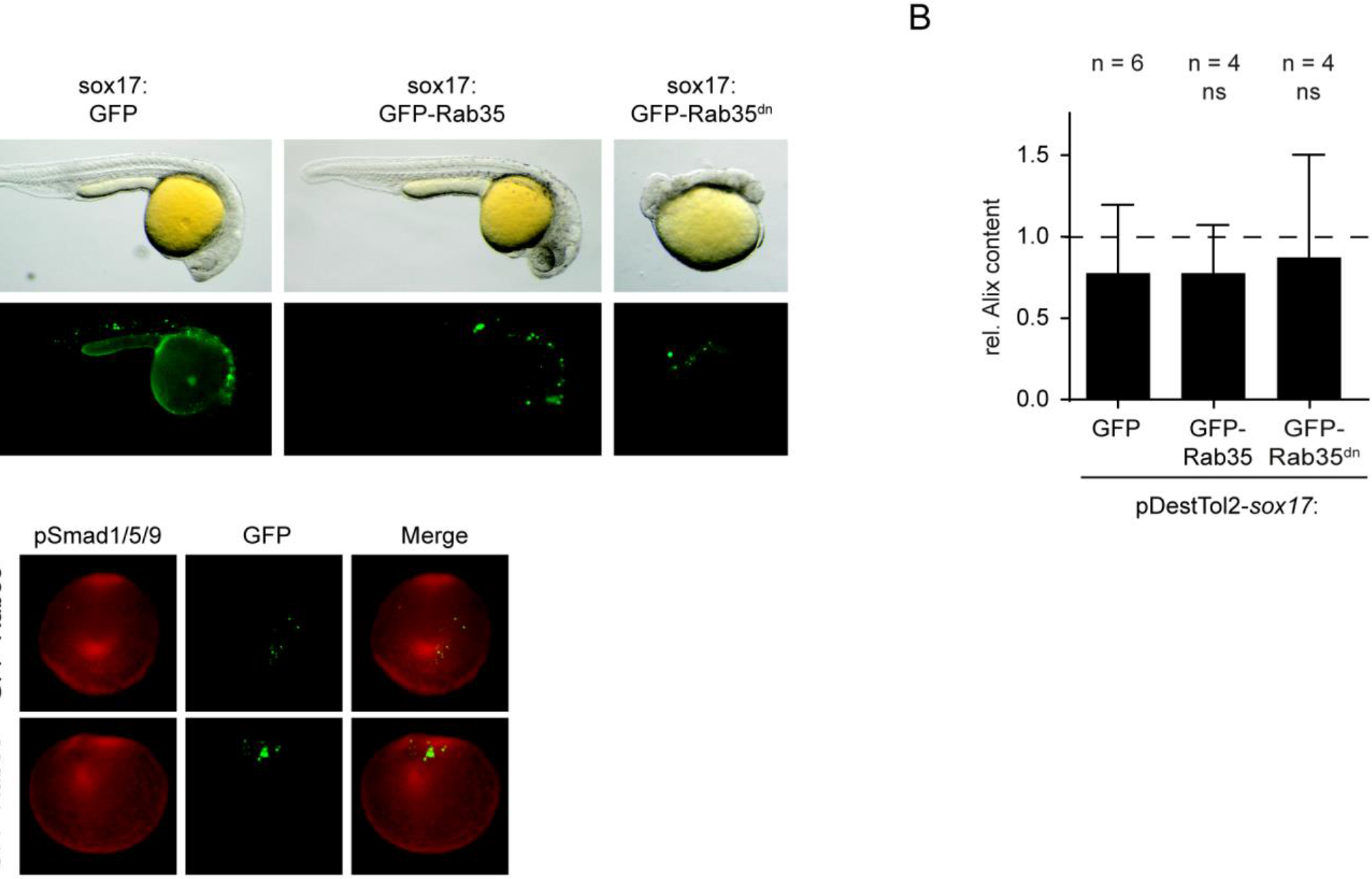
Mosaic expression of dominant negative Rab35 in the endoderm does not result in a measurable reduction in secreted EVs. (A) Images of zebrafish embryos at 24 hpf expressing either pDestTol2-*sox17*:GFP, pDestTol2-*sox17*:GFP-Rab35, pDestTol2-*sox17*:GFP-Rab35^dn^. (B) The amount of secreted EVs in transgene-expressing zebrafish was measured by determining the ALIX-content in EV-isolates of 10 hpf-zebrafish embryos using western blotting. The values were normalized to the wildtype-control and the means were compared using ANOVA and Tukey’s post hoc test. (C) Representative images showing the merge of the GFP-fluorescence representing the transgene and pSmad1/5/9-immuno fluorescence staining.

**Figure S2.**
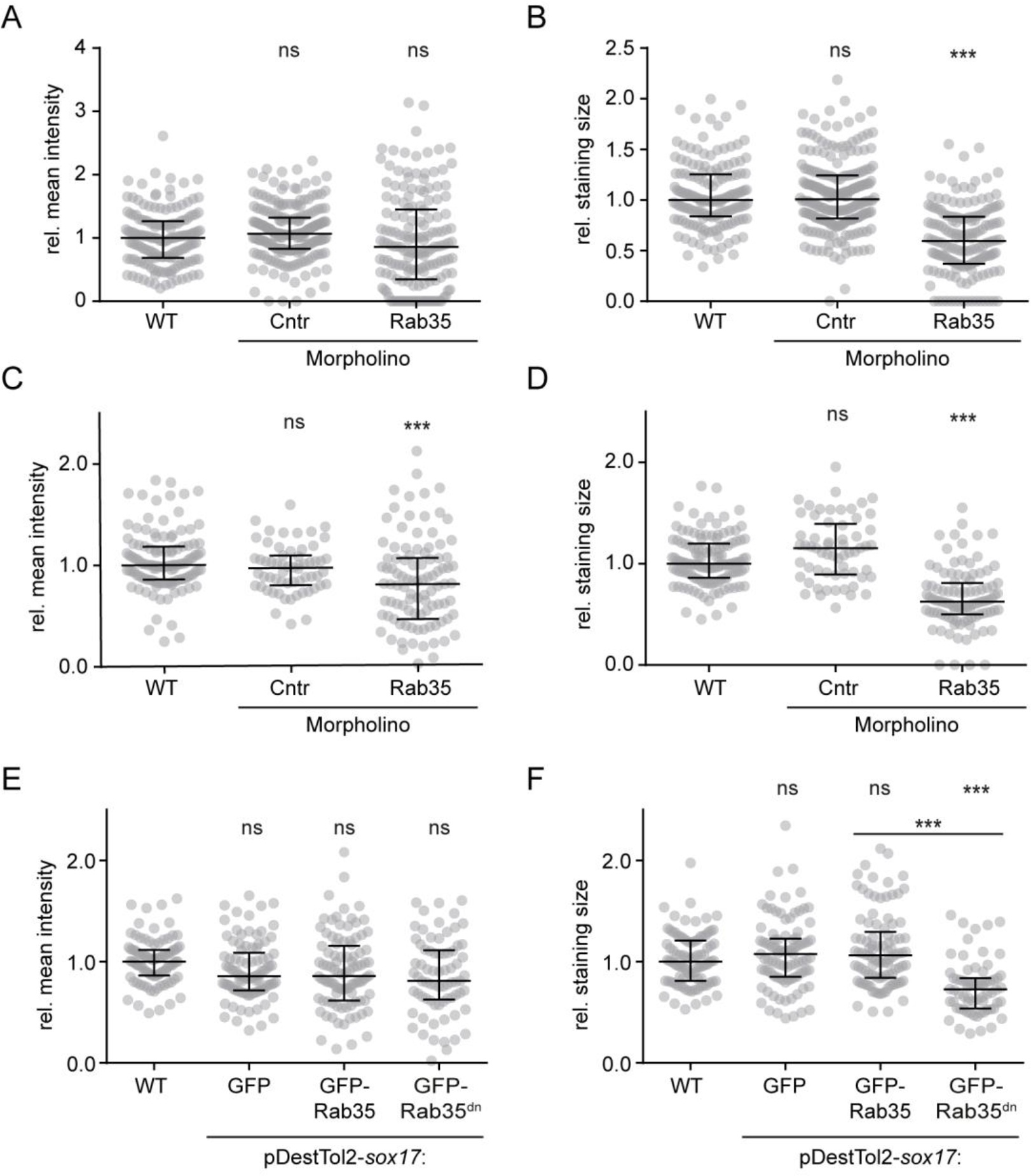
The staining area is the predominant factor determining the reduced *in vivo*BMP-signalling response after Inhibition of EV-secretion. Scatter-whisker-charts plotting the parameters used to calculate the *nkx2.5*-expression A, B; Figure 3B) and total fluorescence in the tail buds of pSmad1/5/9 immunofluorescence stainings (C, D; Figure 3D) in Rab35-morphants as well as the total fluorescence in pSmad1/5/9- immunofluorescence stainings of zebrafish embryos expressing GFP-Rab35^dn^ in the endoderm (E, F; Figure 5D). (A, C, E) The background-corrected mean intensity measured in the labelled regions of interest. (B, D, F) The area of the regions of interest normalized with the area of the embryo. Sample distributions were compared using the Kruskal-Wallis-test with Dunn’s post hoc test.

